# Brain-Guided Convolutional Neural Networks Reveal Task-Specific Representations in Scene Processing

**DOI:** 10.1101/2025.01.02.631092

**Authors:** Bruce C. Hansen, Michelle R. Greene, Henry A.S. Lewinsohn, Audrey E. Kris, Sophie Smyth, Binghui Tang

**Affiliations:** Department of Psychological & Brain Sciences, Neuroscience Program, Colgate University, Hamilton NY; Barnard College, Columbia University, Department of Psychology, New York, NY

**Keywords:** Scene understanding, electroencephalography (EEG), convolutional neural networks (CNN), brain-guided neural networks

## Abstract

Scene categorization is the dominant proxy for visual understanding, yet humans can perform a large number of visual tasks within any scene. Consequently, we know little about how different tasks change how the scene is processed, represented, and its features ultimately used. Here, we developed a novel brain-guided convolutional neural network (CNN) where each convolutional layer was separately guided by neural responses taken at different time points while observers performed two different tasks on the same set of images. We then reconstructed each layer’s activation maps via deconvolution to spatially assess how different features were used as a function of task. The brain-guided CNN made use of image features that human observers identified as being crucial to complete each task starting around 244 ms and persisted to 402 ms. Critically, because the same images were used across the two tasks, the CNN could only succeed if the neural data captured task-relevant differences. Our analyses of the activation maps across layers revealed that the brain’s spatiotemporal representation of local image features evolves systematically over time. This underscores how distinct image features emerge at different stages of processing, shaped by the observer’s goals and behavioral context.

## Introduction

The human brain rapidly encodes and represents complex scene information from our visual environment, enabling a variety of intelligent behaviors. A leading framework for characterizing visual representations of scenes is a hierarchical feedforward sweep through the ventral visual system. In this process, features of increasing complexity are extracted and combined to achieve a semantic classification of the environment [1-6]. This framework has been supported by psychophysical (e.g., [7]), MEG (e.g., [8]), EEG (e.g., [9-10]), and fMRI (e.g., [11-12]) evidence. It has also inspired the development of deep convolutional neural networks (CNNs) in computer vision, which can achieve human-level classification performance [13], and are known to correlate with different processing stages (both spatially and temporally) along the visual hierarchy (e.g., [14-15]).

Despite these successes, the feedforward model relies on two oversimplifications that limit our understanding of scene representations. First, although scene categorization is an important aspect of visual perception (e.g., [16]), it is unlikely to be the only goal. We use different information in an environment to achieve different goals. For example, walking through an environment may only require understanding major paths and their relative difficulty (e.g., [18-19]), while searching for a specific object engages knowledge of what the object looks like and where it is likely to appear (e.g., [20-23]). Second, neural scene representations are highly dynamic, even with static images (reviewed in [24]). This suggests that the underlying representational codes for scenes are not fixed but vary spatially (across the scene) over time—even in the earliest stages of processing [25-26]. Therefore, a more complete characterization of complex visual processing should examine how the visual system uses specific scene features over time and how this varies with the observer’s task.

In previous work, we have shown that neural responses to images of scenes use different features over time and that an observer’s task can dramatically alter feature use [9]. For example, relatively low-level features, such as multi-scale oriented filters, were only used when observers discriminated the dominant orientations in a scene. In contrast, the lexical distance between category names was only used when observers assessed the similarity between scene category names. This suggests that the brain does not simply create a fixed categorical state based exclusively on the environment. Instead, it creates a representation that best matches the behavioral goals of the observer, likely resulting from a dynamic interplay between expectations and attention [27-32]. While that approach revealed how global task-modulated scene information was used over time, it did not examine how task-related information varied over image location, nor how different image locations were prioritized dynamically over time.

To address this issue, Hansen and colleagues [26] developed a spatial mapping analysis for EEG data called Dynamic Electrode-to-Image (DETI) mapping. This method provided a much more refined understanding of the neural dynamics for scenes and established when different scene regions contribute to the neural code. Specifically, DETI mapping established how spatial frequency-selective neural populations prioritized different image locations over time. However, the mapping of EEG responses to image regions was only carried out with one experimental task, so we could not determine how an observer’s task altered the image-region coding of scene information.

Ideally, we would map the image locations containing task-relevant information to real-time neural signals. While DETI showed that this is straightforward for easily parameterized features such as spatial frequency, scaling this approach to the higher-level representations underlying goals and tasks is not trivial. A major hurdle is establishing the utility of each image region for any given task, which involves numerous assumptions about what should be considered useful and when it should become useful.

As CNNs link image location information with higher-level outputs, this work investigates training a novel CNN that processes images as input and uses the corresponding neural responses over time to guide how the weighted image information is passed through each subsequent layer. Previous studies have used “brain guidance” to improve CNN robustness for classification by aligning CNNs with neural responses [33-42]. However, our approach visualizes the image features used while participants perform different visual tasks. Essentially, this involves training a CNN to differentiate between identical images using neural signals from participants performing different tasks on those images. Stated differently, we asked the network to take in an image and predict what task the observer was engaged in while viewing that image. CNNs require structural differences between images to differentiate them. Without neural guidance, a CNN would fail to differentiate between identical sets of images. Therefore, our approach forces the network to evaluate each layer’s filter responses against task-evoked neural responses to achieve accurate task classification.

This brain-guided CNN approach allows us to reconstruct activation maps (via deconvolution [43]) for each CNN layer and image. Those maps can illustrate the image regions containing task-relevant information (as measured by EEG) at a given time point in visual processing. We then compare those maps against maps generated from human behavior to show when and how different image features may support task-oriented behavior. Critically, our aim is not to improve CNNs per se, but to use a brain-guided CNN as a tool for understanding how and when the visual system uses task-relevant features. Our results show that a CNN can successfully differentiate between identical sets of images when augmented with neural responses sampled at different timepoints. Crucially, the network makes use of task-relevant image regions beginning around 244 ms to achieve successful task classification. Interestingly, the network struggled to classify the task for images where the task-relevant regions were largely overlapping. This indicates that neural responses generated by those images were more similar between the two tasks, which would be expected if task-relevant information were accurately captured by the CNN. This study thus advances our understanding of how the visual system prioritizes task-relevant visual information over time.

## Results

Participants (N = 24) completed two pre-cued tasks (object and scene function) on the same set of images while their neural responses were recorded via EEG, generating two separate ERP datasets (one per task). The object task assessed participants’ confidence in whether a particular cued object was present in the image, while the function task evaluated the likelihood that a specific cued action could be performed in the scene (see the Methods section for further details). Additionally, an independent set of observers (N = 50) viewed the same image set and marked image regions they believed contained useful features for each task. Each image was marked separately for the object and function tasks. The behavioral experiment generated task-relevant ‘information utility’ (IU) maps for each image, identifying regions relevant to each task. IU maps were used to assess the extent to which each layer in our brain-guided CNN utilized the same regions as human participants for performing each task. Only the participant averaged ERP data for each task were used to guide the CNN during training. After a brief discussion of our behavioral results (Section 1) and introductory remarks on the brain-guided CNN (Section 2), we show findings from our analyses of the convolutional layers in that network (Sections 3 - 4).

### 1.0 Overall Behavioral Responses

We constructed information utility (IU) maps to provide a behavioral reference for analyzing the image reconstructions at each network layer (detailed in the next section). Those maps were built based on 50 independent observers who each clicked on 10 image regions that they thought were the most relevant to either the possible locations of an object, or locations relevant for a scene function. Example IU maps, along with the averages across all maps within each task, are shown in **Figure 1**.

**Figure 1.**
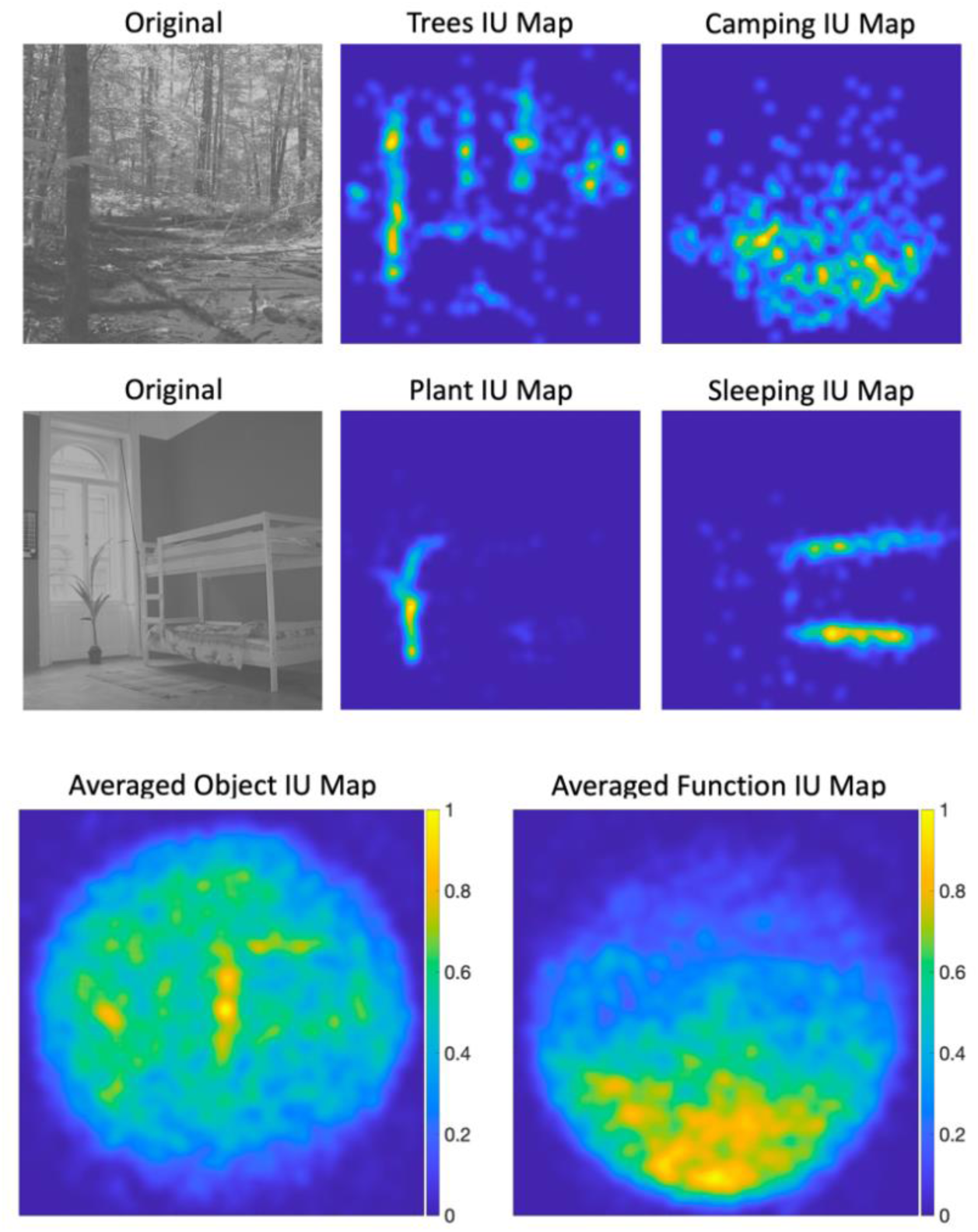
Top two rows: example object and function IU maps for two different stimuli. Bottom row: averaged object and function IU maps (averaged over all stimuli). See text for further detail.

The averaged maps in **Figure 1** show that relevant locations for the object task were distributed broadly across image space, with a slight concentration in the center. For the function task however, the most informative regions were largely concentrated along lower portion of the image. In subsequent analyses, we examine the extent to which the brain-guided convolutional neural network’s layers correspond to these task-relevant regions.

### 2.0 The Brain-Guided Convolutional Neural Network

The basic architecture and general operations of the brain-guided convolutional neural network presented here are inspired by those of AlexNet [44], see a detailed account of the brain-guided architecture in the Methods section, and **Figure 2** for an illustration. Each network layer was guided by a different ERP timepoint that represented times where neural responses were maximally dissimilar across tasks (see Methods section). Neural dissimilarity between tasks is important because it suggests task-based information is being encoded. Further, because the neural data are the only means for the network to differentiate between identical images, it is important that the network is guided by neural responses that are as distinct as possible across tasks.

**Figure 2.**
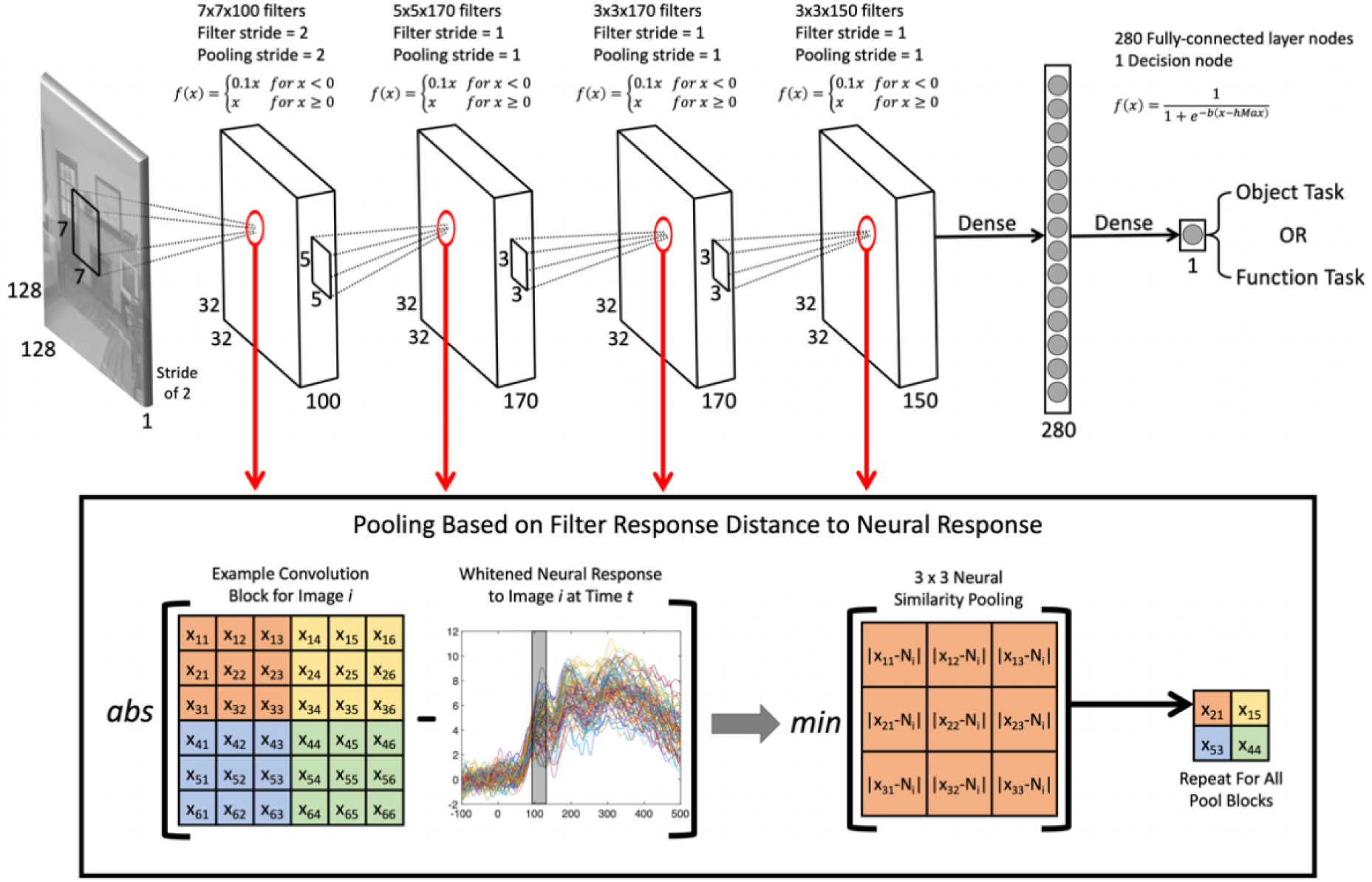
Illustration of our brain-guided CNN and neural similarity pooling operation. Top: the network consisted of four convolutional layers, each being guided by neural responses sampled from one of four different sequential time points from ERP data measured across all images and both tasks. The output of the network consisted of a single decision node that predicted which task an observer was performing on the image. Bottom: we replaced the typical max-pooling operation with a neural similarity pooling operation that selected filter outputs that most closely matched the neural response elicited by each image (see Methods section for further detail).

The network was trained on the same 64 images used in the EEG and behavioral experiments (see Methods for more detail about the images). During training, the network predicted a given image’s task using the image’s pixel values and its corresponding task-driven neural responses at four different time points (one for each convolutional layer). Once trained, we used a deconvolution algorithm [43] to reconstruct the activation maps for each input image and each layer and analyzed those maps against the IU maps in order to identify how and when task-related features become available.

### 3.0 Network Performance and Analysis Results

We trained our brain-guided CNN 10 times, each time using a different random initialization of the weights across all layers. The following sub-sections detail the results of the model’s performance analysis, as well as our behavioral response and reconstruction analysis.

### 3.1 Network Classification Performance

Across the 10 training runs, the brain-guided CNN correctly predicted the task that the observer was engaged in 90% of the time. Specifically, we stopped network training once ∼90% accuracy was achieved due to the limited gain in accuracy with more learning epochs. The decision node consisted of a traditional sigmoid activation function, with a correct object task classification being a zero and a correct function task classification being a 1. Object classifications were counted as correct if the sigmoid output fell below 0.4 and function classifications were counted as correct if the sigmoid output was above 0.6. Outputs that fell within the 0.4-0.6 range were counted as incorrect. **Figure 3** shows the response to each image, averaged across training run, when paired with neural responses to the object task (red bars) and to the same set of images when paired with neural responses for the function task (blue bars). The difference between each identical image pair was assessed for statistical significance using a paired samples t-test. All but one of the unique 64 pairs were found to be statistically significant (all P’s < .003). The image that did not reach statistical significance was trending toward significance (P = 0.083).

**Figure 3.**
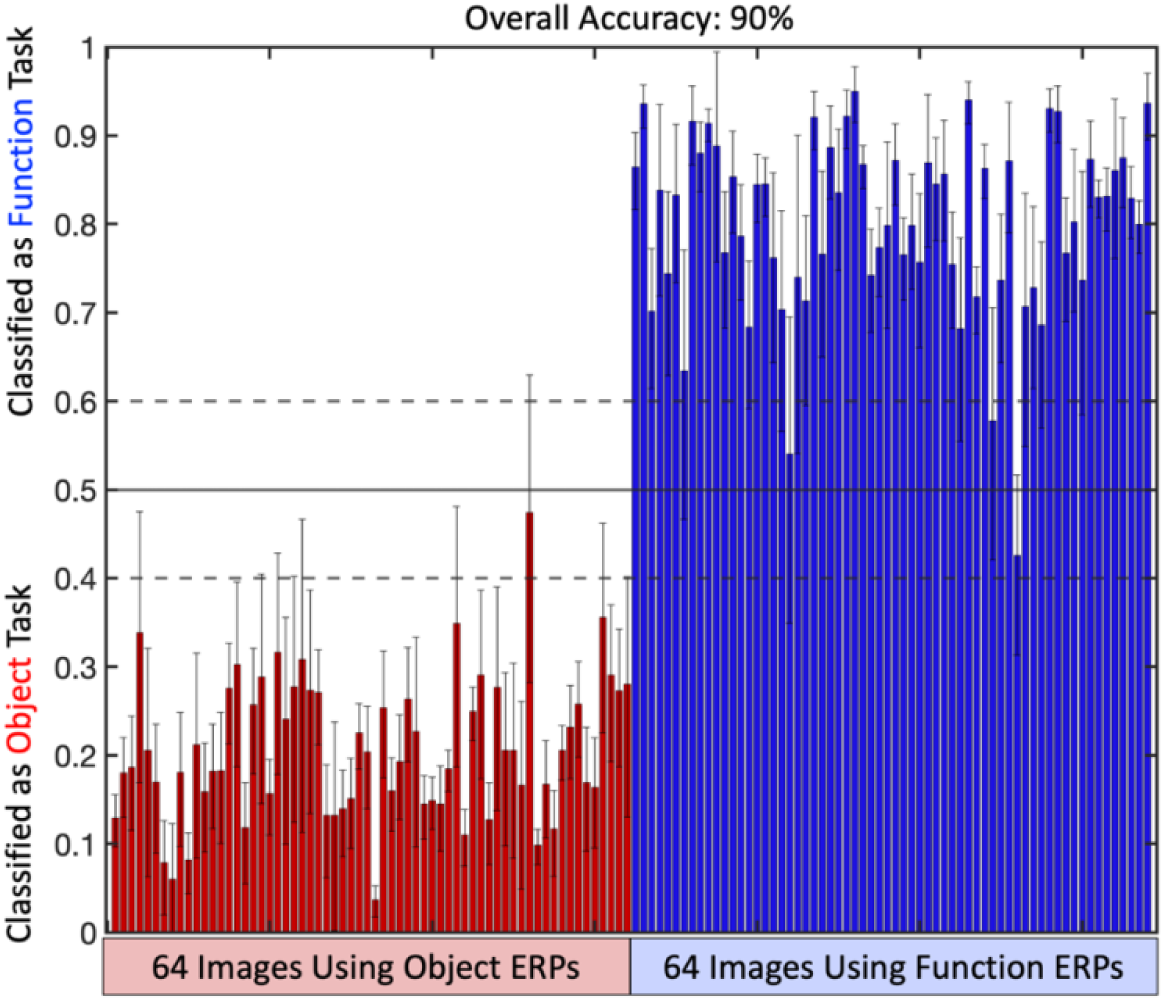
Overall network performance. Image order is identical for the two sets of classifications. Error bars are 95% confidence intervals computed over the 10 different training runs.

### 3.2 Layer-by-layer Stimulus Reconstruction Analyses

Next, we examined the extent to which reconstructed images at each convolutional layer could predict the IU maps that provided task-relevant image locations. This allows us to assess whether and when the network makes use of task-relevant image regions.

For each task, stimuli were reconstructed as activation maps across the convolutional layers using the deconvolution procedure outlined by [43]. Briefly, this process involves passing an image through a trained network and performing the following steps: (1) unpooling the pooled neural similarity output by mapping it back to its original pre-pooling locations, (2) applying the leaky ReLU operator to rectify the responses, and (3) convolving the unpooled maps with the transposed filters. The resulting activation maps were summed and normalized across filters to create a final activation map for each image and task, producing one activation map per image-task pair. **Figure 4** illustrates this processing pipeline. Additionally, activation maps were generated for each of the 10 training runs, resulting in 10 activation maps for each image and task.

**Figure 4.**
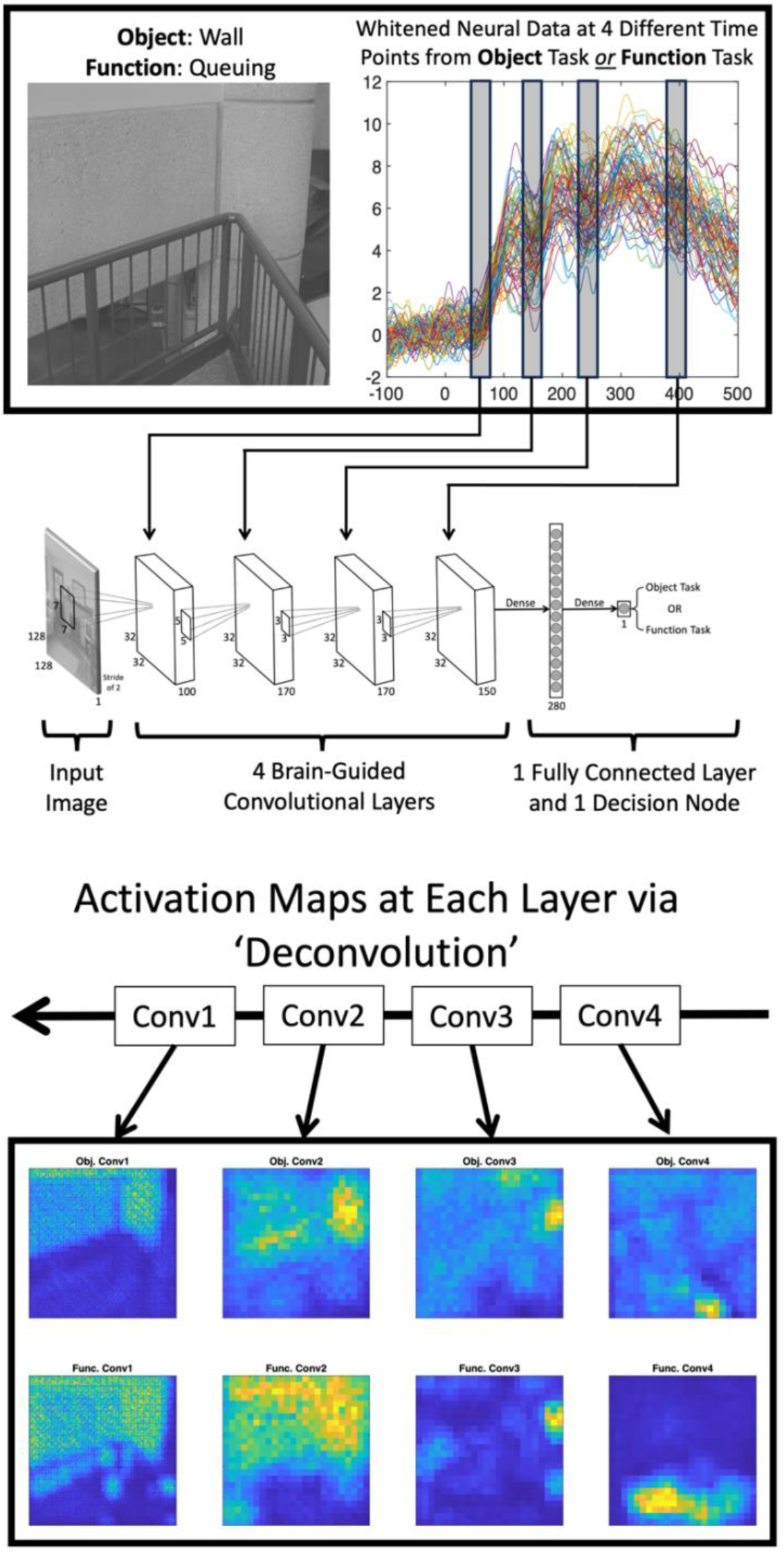
Illustration of the network training and evaluation pipeline for a single image. The network was trained stochastically. Each time an image was passed through the network, it was accompanied by either neural data from the object task or the function task. Once the network was able to classify the task at ∼90% accuracy, we computed activation maps for each layer and each task.

Before conducting the activation map analysis, we estimated a null distribution for the activation maps by creating activation maps for each task whereby the filter weights for each layer were randomized (i.e., untrained weights), and the neural responses were shuffled across images. We repeated this process 50 times with different random weights and a different shuffled order of neural signals for each (50 random activation maps for each layer, image, and task). Next, for each activation map, we computed the dot product between each 3x3 region in each activation map to the corresponding location in each of the random activation maps. We then took the average across all of those distance maps for each image and task. Each of the averaged distance maps were normalized to 1 and then subtracted from 1. That ensures that regions with high dissimilarity (where the activation maps contained activations that were much larger than expected by chance) had the highest values. The dissimilarity maps were then used to weight the activations maps, thereby ensuring that only activations that were greater than expected by chance would carry more weight in the activation map analysis.

To compare activation maps between the two tasks at each layer, we first took the difference between a given object activation map and function activation map. At locations where the differences were greater than zero (larger activation for objects than functions), the difference value was mapped to the corresponding location in a new activation map (an object-only activation map). At locations where the differences were less than zero (larger activation for functions than objects) the absolute value of the difference value was mapped to the corresponding location in a new activation map (a function-only activation map). Each object-only and function only activation map was then weighted by the activation dissimilarity maps mentioned in the previous paragraph. Lastly, the non-normalized dot product was taken between each activation map and it’s corresponding IU map. This process was repeated for each image and task and then averaged over all images within a given task. See **Figure 5a** for an illustration of this analysis. As a relative control, a given activation map was dotted with the IU map from the other task (e.g., dot product between an object activation map and function IU map from the same image). The larger the products between matching activation maps and IU maps relative to mismatched, the more the network was using regions specific to neural response that the task elicited. The above analysis was repeated for each of the 10 training runs and averaged (**Figure 5b**). Independent samples t-tests conducted on each match vs mismatched map pair show that task relevant region information use for both the object and functions tasks begins around 244 ms and persists out to at least 402 ms. Importantly, the relative differences that were observed at 244 ms and 402 ms were observed for each of the 10 training runs and were therefore robust to different random weight initializations.

**Figure 5.**
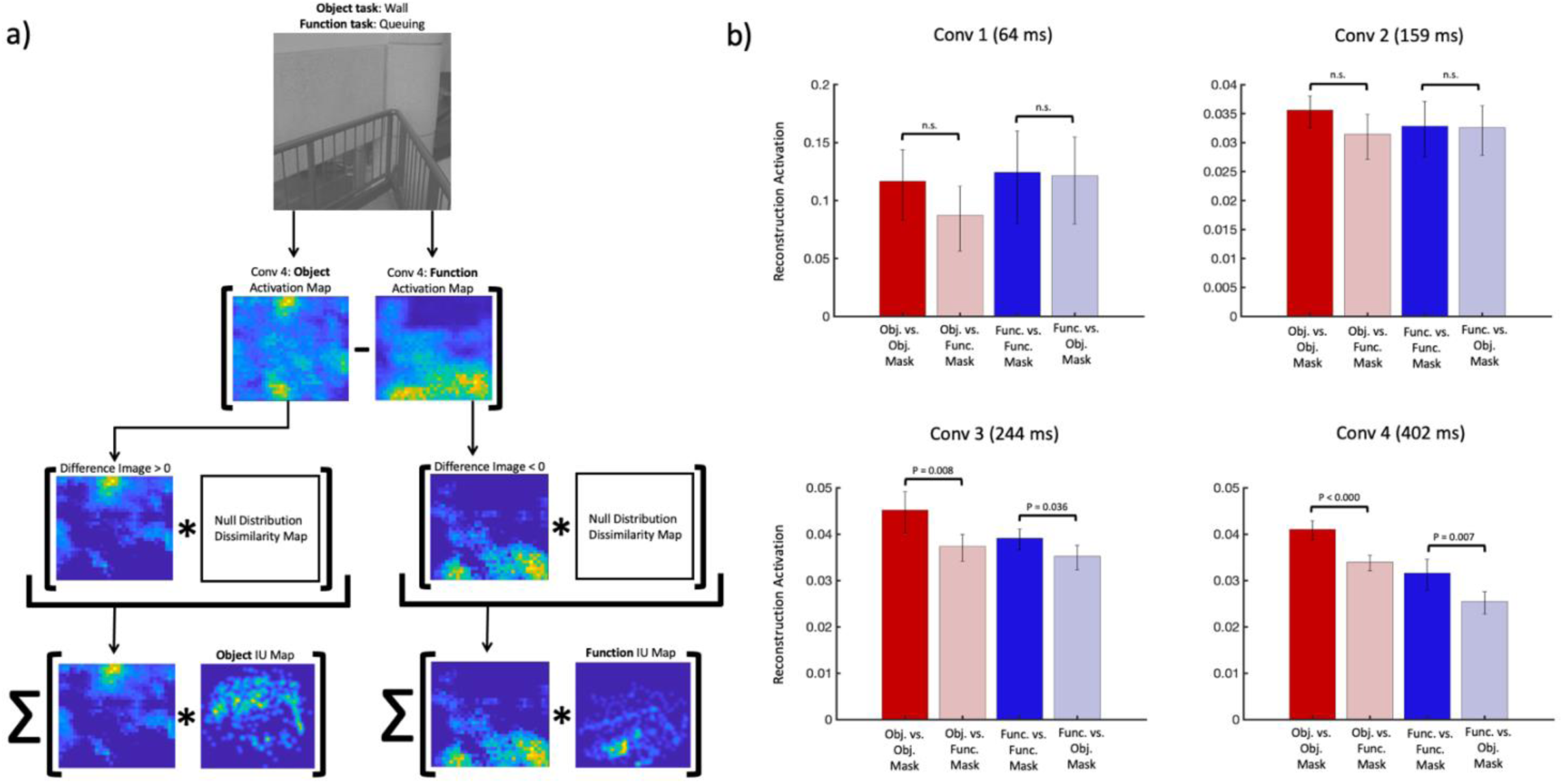
**a)** Illustration of our activation map analysis for an example image and network layer. We first took the absolute difference between two corresponding activation maps for any given stimulus and weighted those values by activation maps that would be expected by chance alone. We then took the dot product between the weighted difference maps and their corresponding task IU map (and the control map from the opposite task). See text for further detail. **b)** Results from our activation map analysis described in the text (and illustrated in Figure 5a). Error bars are 95% confidence intervals computed over 10 training runs. The network is considered to be using task relevant information when the reconstruction activation is higher when the reconstruction activation came from the same task as the IU map compared to when that same activation map was weighted by an IU map from the other task. For example, when tasked with classifying a given image when given neural data from the object tasks, the network used object relevant information when compared to information that was more relevant to the function task.

We next examined the relative dissimilarity (Euclidean distance) of each image’s activation maps across both tasks at each convolutional layer. The paired distances were then normalized and submitted to a one-way ANOVA, which yielded a significant effect of layer, F(3,252) = 7.53, P < .000, with conv1 M = 0.278, conv2 M = 0.363, conv3 M = 0.468, and conv4 M = 0.399. Follow-up independent t-tests revealed significant differences between conv1, conv2, and conv3, (all Ps < 0.03) indicating that as the images made their way through the brain-guided network, the distances between activations for each image between the two tasks were increased, suggesting more distinct task-relevant representations emerged over time.

One question raised by the results shown in **Figure 5b** is whether all four convolutional layers were necessary, or whether a simpler network guided at the relevant time points could achieve a similar result. To test that, we scaled back our network to only have one convolutional layer (each time point used in the full network, separately), followed by a fully connected layer and decision node. We found that, while the smaller network could learn to differentiate the tasks, it did so in a way that did not utilize task-relevant information, suggesting that the representations built up across earlier layers are necessary to create later task-relevant representations. Lastly, to test whether the apparent build-up of task relevant information use is not simply a byproduct of having more layers in the network, we shuffled the time-specific neural data across the layers and conducted the activation map analysis using the behavioral IU maps. The results showed that the network made use of local region information in a manner that was inconsistent with behavior (all Ps > .05 for all match vs mismatched IU map pairs).

### 3.3 Assessing the degree of overlap across tasks

The averaged IU maps in **Figure 1** suggest that, to a reasonable extent, function and object information tended to have different location distributions in our images. Nevertheless, **Figure 6a** also shows signs of overlap between the regions useful for each task. However, the critical question is whether the IU maps (object and function) for each image are sufficiently different. To asses that, we computed the Euclidean distance between each image’s utility maps in MDS space (**Figure 6b**). The distance distribution shows a slight bimodal tendency, suggesting two distributions, one for a set of images with similar IU maps and another with largely dissimilar IU maps.

**Figure 6.**
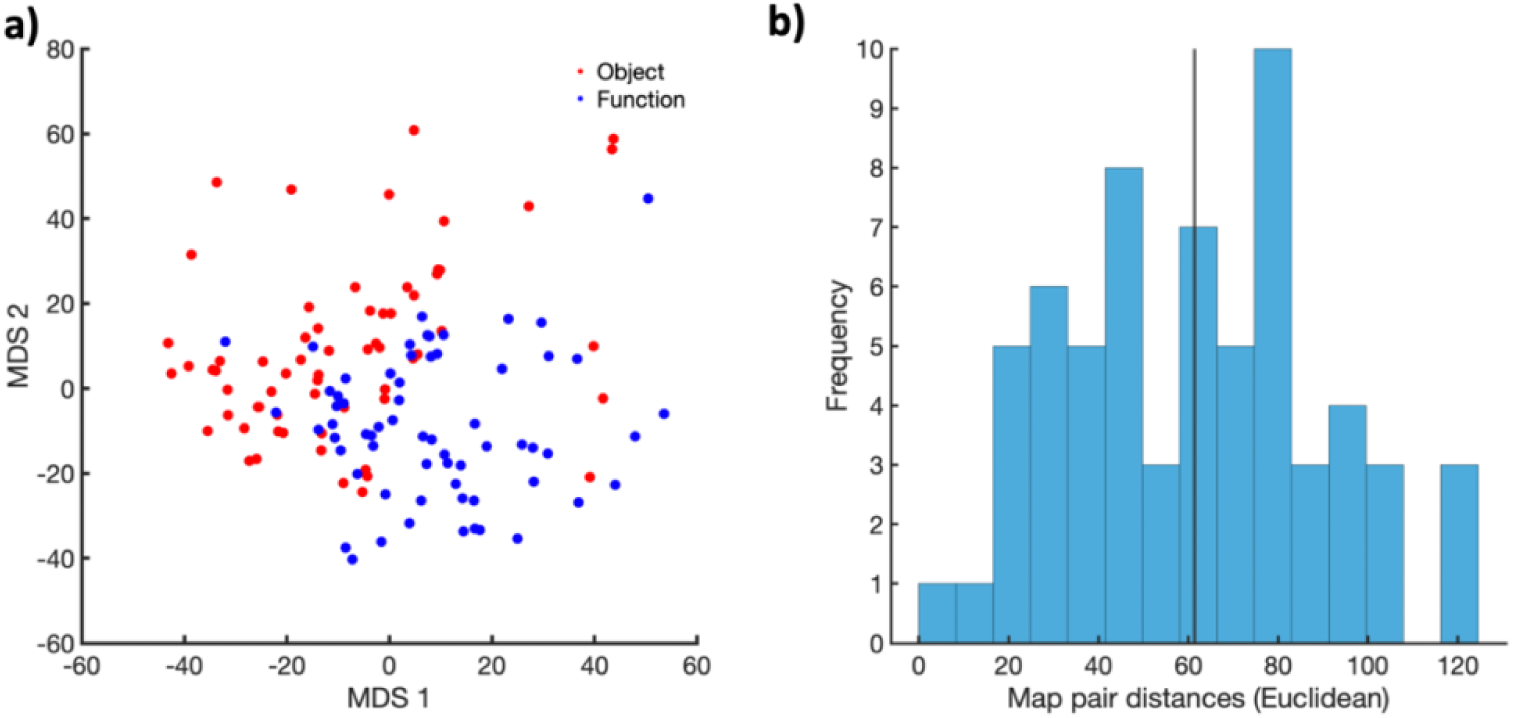
(a) the first two MDS dimensions for the object (red) and function (blue) IU maps. While there is good separation between the types of IU maps, there is some overlap. (b) Euclidean distance for each IU map pair showing an approximately bimodal distribution.

To assess whether the differences between task relevant regions overlap across the images could explain the differences between the ERPs measured for each task, we conducted a split-half analysis along Euclidean distances (x-axis line in **Figure 6b**). That is, the ERP data were split according to images with similar IU maps (N = 32 images) between the two tasks (images below the median) and images with dissimilar IU maps (N = 32 images). We the computed a time-resolved (11ms sliding window) regression analysis between windowed ERPs from each task (**Figure 7**). Larger R2s indicate smaller differences between the ERPs for each task. The R2 plots show overall higher R2s for images with similar IU maps, suggesting that task effects were weaker for those images. By contrast, images with largely dissimilar IU maps had smaller R2s. To assess the extent of the R2 differences as they relate to the timepoints and electrodes that we used for network training, we averaged the R2s across the relevant electrodes within 11 ms windows centered on each of our four target timepoints (**Figure 7**). The overall R2s are relatively low (all less than 0.25) due to our efforts to select time points and corresponding electrodes with low similarity (see Method section), yet there is a trend for increasing dissimilarity over time between the two sets of images.

**Figure 7.**
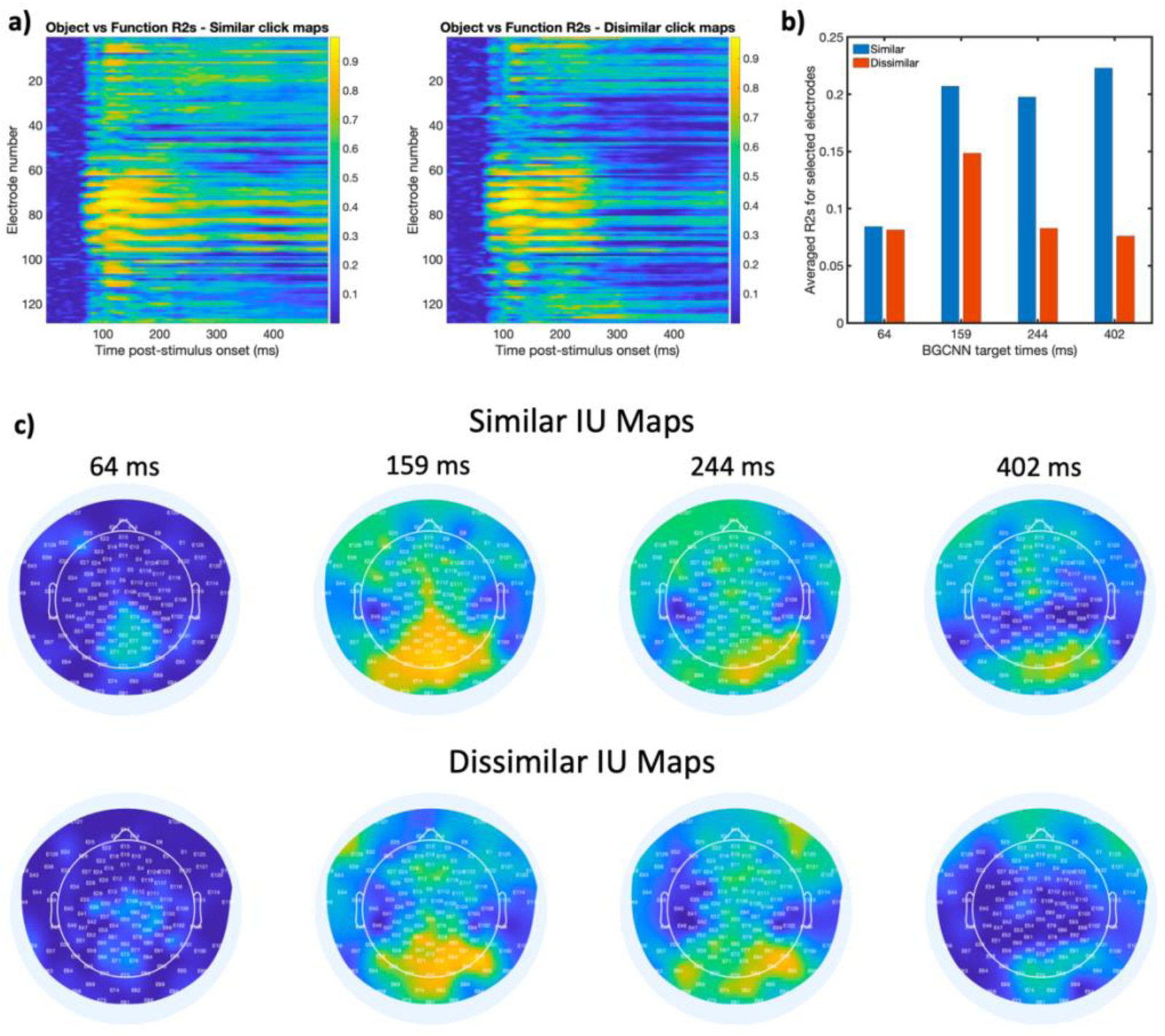
(a) Time by electrode matrices showing time-resolved R2 values between ERP data from each task, but for images with similar IU maps (left) or dissimilar IU maps (right). Lower values indicate the potential for task related ERP effects. (b) Summary plot showing averaged R2s at the targeted time points and electrodes. Note the reduction in mean R2 over time between tasks for images with dissimilar IU maps. (c) Topographic scalp plots showing between task R2 values at the four targeted time points for stimuli with similar (top) vs dissimilar (bottom) IU maps.

Is the brain-guided CNN’s accuracy lower for classifying images with similar IU maps because their ERPs are less distinct and their task-relevant regions overlap? To investigate that, we analyzed the model’s classification performance across 10 training runs. We calculated the difference between the decision node’s output and the target response for each task (0 for objects and 1 for functions), and averaged these differences for images with similar and dissimilar IU maps, and performed an independent samples t-test, which resulted in a significant difference between the two sets, t(62) = 2.15, P = 0.0357, Cohen’s d = 0.54. The mean difference for the similar IU map group M = 0.2231 was larger compared to the dissimilar IU map group M = 0.1806. Smaller values mean better classification, with 95% confidence intervals of [0.002 0.082]. These results suggest that while the model performs well overall, it struggles more with images where the task-relevant regions overlap, likely because those images produce ERPs that were more similar between the two tasks.

One possible explanation for this difference could be that the images themselves differed with respect to their physical characteristics. We examined image-based differences between images that generated similar versus dissimilar IU maps within a tSNE-based image space (see the Methods section for more detail). No apparent clustering of images with similar vs dissimilar IU maps was observed. It therefore does not seem that the differences in ERP similarity between the two tasks can be accounted for by low-level image characteristics (see **Supplementary Figure S1** for more details).

## Discussion

The results of the performance analysis on our brain-guided CNN show that neural responses generated by identical images with two different tasks could be used to guide a relatively standard CNN to differentiate between identical sets of images (**Figure 3).** Crucially, not only could the network learn to differentiate identical images based on task-related neural variance, but it did so by making differential use of image regions that an independent group of raters found useful to complete each task. Further, the network was relatively stable across different learning sessions with different filter weight initializations in terms of both task classification accuracy, as well as the associated activation map constructions obtained via deconvolution. Task-relevant feature use emerged relatively later in neural processing (starting at ∼244 ms, **Figure 5b**). That finding is consistent with previous M/EEG work [45, 9, 46]. For example, our previous work examined when different task-related features begin to explain unique ERP variance [9]. However, that work did not attempt to localize the neural responses to task-relevant regions. Other work has mapped local task-relevant features to neural responses by creating stimuli where task-relevant information is localized only in specific, non-overlapping image locations [45]. While this is a powerful inferential aid, real-world scenes contain regions that are potentially useful across multiple tasks. By contrast, this work addresses those shortcomings by revealing that image locations that support task demands are prioritized in the neural responses.

The analysis of the behavioral IU maps revealed, across all of our task prompts and images, that most of the object-relevant regions of information were broadly distributed with a slight central concentration, while the function-relevant regions were largely concentrated around the bottom of the images. Nevertheless, at the individual map level, we found an approximate bimodal distribution whereby half of our images could be quantified as containing overlapping task relevant regions. That is, our independent raters found certain regions in some of our images useful to complete both the object and the function task. The other half of our images could be classified as having task informative regions that largely did not overlap. Strikingly, this result predicts differential task-related effects. For example, if the neural data are separable based on task, ERPs generated by images with similar IU maps should be more similar compared to ERPs evoked by images with dissimilar IU maps. **Figure 7** shows the results of that analysis and suggests that when testing for task related effects in neural processing, it is important to consider the relative overlap between informative regions across tasks. Further, our image structure analysis (**Supplemental Figure S1**) shows that this relationship cannot be explained by differences in the structural content of the images. Lastly, not only did the relative overlap between informative regions across tasks predict task effects in the ERP data, but it also coincided with the brain-guided network’s performance on images with similar IU maps vs. those with dissimilar IU maps. Specifically, the network struggled more when differentiating identical images that yielded similar IU maps compared to those with dissimilar IU maps. To our knowledge, this is the first investigation to report such a relationship and should be very useful for future work in understanding how behavioral goals in visual environments influence the neural representation of task relevant information.

This work used neural information to guide the training of a CNN. Unlike other work that used brain guidance to improve CNN performance [33-42], our goal was to leverage the localization power of CNNs to provide insight into task-related visual processing differences. However, future computer vision efforts may be aided by understanding how the human visual system uses different scene features for various tasks based on the approach presented here.

While the success of our brain-guided CNN opens new and exciting paths for similar applications in other domains of systems neuroscience, it is not without its limitations. For example, the neural similarity pooling operation that we used here relies on finding the smallest distance between an image’s neural response and filter outputs within each pooling region. This can be problematic for noisier datasets where the pattern of neural responses across images is less stable, thereby making it harder for the network to differentiate images based on task in any meaningful way. One possible solution would be to incorporate a fitting procedure within the similarity pooling operation that considers how any given filter responds across all images in the stimulus set, and then maps the residual between the input image’s response and the fit to the corresponding location within a given pooling region. Then, as with our current neural similarity pooling operation, find the smallest residual in the pooling region and map its corresponding filter response to the pooled output image for that filter. Because the fitting would be relative to all images, the residual would be more stable for each training iteration. We did pilot that approach with a single brain-guided CNN layer, and found that the resulting activation maps had less noise and had more gradual variations between regions of low and high activation. However, because that process requires passing all images through the layer before the fitting can take place, the compute time grew exponentially as more layers were added. Another limitation is that, by feeding each layer’s output into the next, we are assuming that the neural sources at time *x* feed directly (and entirely) into the sources at the next timepoint *x* + Δ*t*. Future developments of this network might handle that by taking into account predictive signaling between electrodes at different time points (e.g., Granger causality) to not only select which electrodes to use for brain guidance, but to also facilitate which (and how many) time points (i.e., convolutional layers) to use. Next, the network was only trained to differentiate between two tasks, so a more robust test of this type of network would include more tasks using the same set of images. Lastly, more convolutional and/or fully connected layers might provide more refined insight into how and when observers make use of task relevant information. It’s also possible that further insights could be gained by using the neural responses to guide network operations other than at the pooling stage. Nevertheless, the network reported here was highly successful in differentiating images based on task-driven neural data, and crucially, made use of the same image regions that human observers found useful to complete the task.

## Methods

### Apparatus

All stimuli were presented on a 23.6” VIEWPixx/EEG scanning LED-backlight LCD monitor with 1 ms black-to-white pixel response time. Maximum luminance output of the display was 100 cd/m^2^, with a frame rate of 120 Hz and resolution of 1920 x 1080 pixels. Single pixels subtended .0382° of visual angle as viewed from 35 cm. Head position was maintained with an Applied Science Laboratories (ASL) chin rest.

### Participants

A total of 36 participants were recruited for this experiment. Of those, nine failed to complete all four recording sessions and three were excluded for having fewer than 50% valid trials following artifact rejection. The age of the remaining 24 participants (17 female, 22 right-handed) ranged from 18-22 (median age = 20). All participants had normal (or corrected to normal) vision as determined by standard ETDRS acuity charts and were compensated for their time. This study was reviewed and approved by Colgate University’s Institutional Review Board, and all methods involving human participants were performed in accordance with the guidelines set forth by that board. All participants gave written informed consent before participating.

### Stimuli

Unless noted otherwise, images were selected using the same methods reported in [26]. Specifically, we sampled images from a large database of real-world scenes consisting of 2500 photographs that varied in content from purely natural to purely carpentered (both indoor and outdoor), with various mixtures of natural/carpentered environments in between [47]. All images were 512 x 512 pixels (subtending 19.55° visual angle) and converted to grayscale using the standard weighted sum conversion in MatLab.

For the purposes of stimulus presentation and analysis, all images were calibrated according to the following procedures. First, each image was fit with a hard-edge circular window (with a diameter of 512 pixels) whereby all pixels that fell outside of the circular window were set to zero. Next, each image was converted to an array, I(y), that included only the pixels that fell within the circular window and were made to possess the same root mean square (RMS) contrast and mean pixel luminance. Holding RMS contrast and mean luminance constant is an important step as it ensures that the variation in neural response magnitude across images is due to structural/semantic variations as opposed to global contrast or luminance variations across the images.

Root mean square contrast is defined as the standard deviation of all pixel luminance values divided by the mean of all pixel luminance values. Image arrays were set to have the same RMS contrast and zero mean using the following operations.

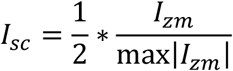

with I_zm_ defined as:

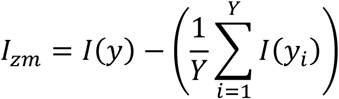

The pixels values of each image array are first normalized to fall between [-.5 .5] with zero mean as follows,

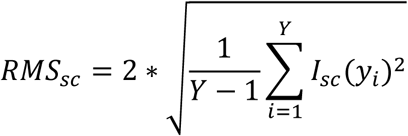

We then calculated an RMS scaling factor, S_rms_ = (2*RMS_t_)/RMS_sc_, with RMS_t_ set to a reasonable target RMS value. By reasonable, we mean a value that did not result in significant (> 5%) clipping of the resulting pixel values. That value was 0.20 for the images used in the current study. Finally, each image array was scaled to have an RMS equal to RMS_t_ and reassign to I(y) as follows: I(y) = 127*(I_sc_*S_rms_). Note that scaling by 127 puts the scaled pixel values of I(y) back in the original range of I_zm_.

Stimulus images were selected according to an image state-space sampling procedure as follows [26]. All images in the database were left in vector form after RMS normalization. When in array form, each cell constitutes a coordinate in a high-dimensional image state-space where each coordinate takes on a pixel luminance value ranging from [-127, 127]. An angular distance matrix was then constructed by calculating the angular distance (in degrees) between each image array and every other image array as follows:

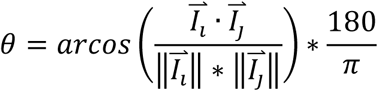

From there, the angular distance matrix was projected into a lower dimensional space (2D) via t-distributed stochastic neighbor embedding (t-SNE) [48]. Projecting the images into this space enables a general lower dimensional organization of images based on their structural attributes defined by pixel luminance.

From this set, 80 images were selected by uniformly sampling from the lower dimensional t-SNE space. Each of the three dimensions was divided into 10 bins, and when possible, one image was randomly selected from each bin. This procedure ensured that different regions in our image state-space were represented in the stimulus set. We chose to use 80 images to have as many images as possible while keeping the overall length of the recording sessions to a practical limit. All selected images maintained their RMS of .20 (defined above) but had their mean pixel luminance set to 127 and then fit with a circular linear edge-ramped window (512-pixel diameter, ramped to the mean pixel luminance) to obscure the square frame of the images. That step ensures that the contrast changes at the boundaries of the image were not biased to any particular orientation [49].

### Assigning Task Targets to Stimuli

Each stimulus image was assigned one object target as well as one function target from a set of 16 functions and 15 objects. The assignments were fixed for each image and did not change across the entire experiment. Assignments were determined by a preliminary experiment that asked observers (N=475, recruited on Amazon’s Mechanical Turk) to check off objects that could be found in a scene, or functions that could be performed in the scene from sets of 17 objects and functions, derived from the Scene Attribute Dataset [50]. One hundred participants ranked each image for both objects and functions, and the most consistently-selected object and function were used in subsequent experiments.

### Experimental Procedure

The current study consisted of two behavioral tasks, namely an object identification confidence task and a function likelihood assessment task. Both tasks were run using the same set of 80 images. The experiment consisted of four recording sessions, with sessions blocked by task (two sessions for each behavioral task). The order of the task blocks was counter-balanced in an alternating sequence across participants (that is, participants never completed two identical task blocks back-to-back). The time to complete each session ranged between 58-70 min, and the time between sessions varied between two days and one week. Within each session, all 80 stimuli were presented 15 times, resulting in a total of 30 presentations per image over both recording sessions for each task (stimulus presentation order within each task block was randomized).

Each trial began with a 500 ms fixation followed by a task cue screen (1000 ms). For the object task, the prompt took the general form of “How confident are you that there is a X here?”, with X set to a particular object for the up-coming stimulus image. For the function task, the prompt took the general form of “How likely are you to X here?”, with X set to a particular function for the up-coming stimulus image. Twenty percent of the images (N = 16) had distractor labels (objects that were not in the image or functions that could not be performed in the image) as manipulation checks to ensure that the participants made an honest effort to engage with each task). The cue screen was followed by a variable duration (500-750 ms) blank mean luminance screen to allow any fixation-driven activity to dissipate. The blank screen was immediately followed by the stimulus interval (500 ms) that was then followed by a variable 100-250 ms blank mean luminance screen, followed by a response screen. The response screen prompted to give a confidence (object task) or likelihood (function task) rating on a 1-4 scale (1 being least, four being most) using a button box. Response time was unlimited, but participants were encouraged to respond as quickly as they could.

### EEG Recording and Processing

All continuous EEGs were recorded in a Faraday chamber using Electrical Geodesics Incorporated’s (MagStim EGI) Geodesic EEG acquisition system (GES 400). All EEGs were obtained by means of Geodesic Hydrocel sensor nets consisting of a dense array of 128 channels (electrolytic sponges). The online reference was at the vertex (Cz), and the impedances were maintained below 50 kΩ (EGI amplifiers are high-impedance). All EEG signals were amplified and sampled at 1000 Hz. The digitized EEG waveforms were first highpass filtered at a 0.1 Hz cut-off frequency to remove the DC offset, and then lowpass filtered at a 45 Hz cutoff frequency to eliminate 60 Hz line noise.

Continuous EEGs were divided into 600 ms epochs, i.e., 99 ms before stimulus onset and 500 ms following stimulus onset (0 ms). Trials that contained eye movements or eye blinks during data epochs were excluded from analysis via magnitude thresholding followed by visual inspection. Additionally, all epochs were subjected to algorithmic artifact rejection whereby voltages exceeding +/- 100 μV or transients greater than +/- 100 μV were omitted from further analysis. These trial rejection routines resulted in a median of 19% (range 4% - 28%) of total trials being rejected across participants for the object task, and a median of 17% (range 2% - 34%) of total trials being rejected across participants for the function task. The difference in rejection percentages between the two tasks was not statistically significant (paired samples t-test, t(23) = 0.372, P = 0.7133). Each epoch was then re-referenced offline to the net average, and baseline-corrected to the last 99 ms of the blank interval that preceded the image interval. Finally, ERPs were constructed for each participant by averaging the processed epochs across trials for each valid image at each electrode, resulting in two 128 x 600 x 64 ERP data matrices for each participant (one matrix per task). Valid images were those with a task cue that was congruent with the content in the stimulus images – distractor images did not contain any useful regional information because their contents did not match either of the task cues. The full dataset (59 GB) is available upon request from the corresponding author.

### Electrode and Time Point Selection for Brain-Guidance

The primary task of the brain-guided CNN was to differentiate between identical sets of images based on their neural responses across the two EEG experiment tasks. It is therefore important that the neural responses from any given electrode and time point for each task be as uncorrelated as possible. That is, it is crucial to select data that is showing a task-based difference in the neural response, otherwise, the network would use near identical neural responses from each task to differentiate between identical sets of images, thereby setting up the network to fail. To identify which electrodes and time points showed significant task-related differences, we conducted a time-resolved R^2^ analysis, comparing participant averaged ERPs from the function task to those from the object task on a per-electrode basis to identify time points and electrodes that differed between tasks. Starting from the stimulus onset and advancing in 1 ms steps, we used an 11 ms time window centered on each time point. For each image and task, we calculated the average ERP within the window, producing two 64 x 1 vectors (one for each task). We then computed the R^2^ between these vectors, then converted it to a dissimilarity using 1-Pearson *ρ*^2^ defined below, with subscript *e* denoting the electrode, *i* indicating a given stimulus, and *μ* being the mean across all images for each task.

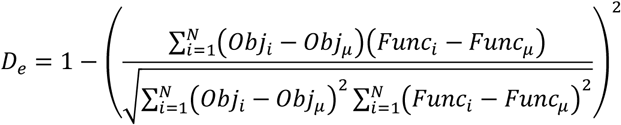

We chose an 11 ms time window so that peak component variability over small and/or short time intervals could be captured. Also, we excluded the electrodes immediately around the ears, on the brow line of the forehead, and cheeks as those electrodes are most susceptible to noise that, leaving 105 electrodes for the neural dissimilarity analysis. After calculating a dissimilarity metric, *D*, for each electrode at each time point, we set a dissimilarity threshold of 0.7, selecting all electrodes with *D*s exceeding that value. That threshold was chosen to ensure that the sphering operation applied to the neural data (explained in the next section) would provide sphered components that closely resembled the original neural data, as sphering highly correlated variables often results in sphered components that do not correlate well with the original variables. Once the electrodes were gathered for each time point, we computed the average *D* across these electrodes on a time point-by-time point basis. We then selected the time points (and corresponding electrodes) with the highest dissimilarity that were separated by at least 100 ms. This separation was chosen to capture distinct ERP signals while still covering the full stimulus interval, avoiding the temporal correlation typical of nearby time points (see **Figure 8** below). Three time points stand out, namely 144 ms (3 electrodes), 244 ms (12 electrodes), and 402 ms (60 electrodes). Unfortunately, the dissimilarity peak at 144 ms only had 3 electrodes, so we used the next peak (159 ms), which had 8 electrodes. All of those time points are relatively late in visual processing, so we selected 64 ms (95 electrodes) as an example of an early time point that is consistent with the initial processing of visual information [51].

**Figure 8.**
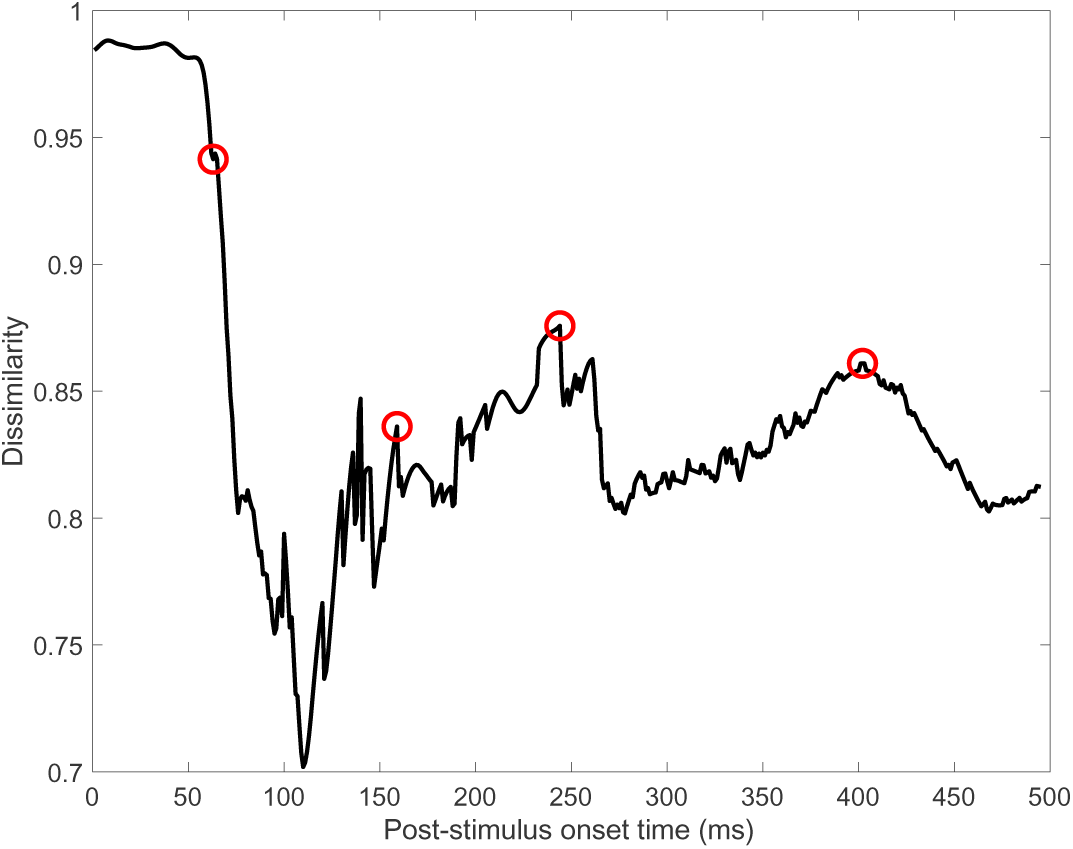
Dissimilarity analysis results.

### Preparing the Neural Data for Network Guidance

Neural data for each task were selected at four time points (and corresponding electrodes) by averaging each image’s ERP within an 11 ms time window centered on each time point on an electrode-by-electrode basis and then averaged over electrodes to yield one neural response per image. This resulted in two 64 x 4 neural response matrices (one for each task), that were then normalized to the absolute maximum at each time point and within each task. To remove any shared variance between and within tasks and between time points, we sphered the data according to all possible task and time point pairs using the zero-phase component algorithm (ZCA, [52]). All sphered components correlated with their original versions at or above r = 0.90. It is the sphered components that were used to guide network operations at each convolutional layer (i.e., one sphered component per image and per task and time point).

### Brain-Guided Convolutional Layers

To ease the computational demand on the network, input images were downsampled from 512 x 512 pixels to 128 x 128 pixels, giving an input layer dimensionality of 128 x 128 x 1. To assist in protecting against exploding gradients, the pixel luminance values of the downsampled images were scaled to 1% of their original RMS-contrast normalized range and set to have a mean pixel value of zero. The first convolutional layer received neural responses from the earliest time point (64 ms post-stimulus onset) and consisted of 100 7 x 7 filters (7 x 7 x 100). Each filter was convolved with the input image with a stride of two pixels and a pooling stride of two pixels, resulting in a 32 x 32 response output for each filter (32 x 32 x 100). The filter dimensions and stride lengths were chosen to aid in constructing well-sampled activation maps (discussed in the Results section) [43]. In terms of spatial frequency, a 7 x 7 filter has a frequency range of 18.28 – 36.57 cycles per picture (0.94 – 1.88 cycles per degree when scaled up to the ERP experiment viewing conditions). Brain guidance was incorporated by replacing the traditional max-pooling operation with a ‘neural-similarity pooling’ operation, applied to the output of the convolution operation. Specifically, for each filter, the output of the convolution between that filter and the image was mapped to a 64 x 64 filter response matrix. A 3 x 3 pooling region was then moved across that response matrix with a stride of two. For each pooling region, the absolute difference between the z-scored filter responses and the neural response was calculated, creating a 3 x 3 matrix of absolute differences. The smallest distance within each pooling region was selected and assigned to a 32 x 32 pooled output matrix for each filter (see **Figure 2** for an illustration). This approach differs from typical max-pooling, as the neural-similarity pooling operation selects filter responses that most closely match the neural response for each image. That is, it identifies the raw filter response in each pool region that best aligns with the neural response. Before being passed to the next layer, this pooled matrix was passed through a leaky rectified linear unit (leaky ReLu) with a slope coefficient of 0.1. The leaky ReLu was chosen to capture both positive and negative neural responses, as ERPs consist of deflections in both directions.

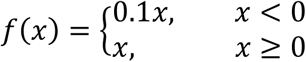

Convolutional layers 2-4 performed the neural-similarity pooling exactly as described for layer 1 (pooling region 3 x 3), with the only exception being that each layer used neural responses from different time points (layer 2 = 159 ms, layer 3 = 244 ms, and layer 4 = 402 ms). Layer 2 consisted of 170 5 x 5 filters (5 x 5 x 170). Each filter was convolved with the output from layer 1 with a stride of one pixel and a pooling stride of one pixel, resulting in a 32 x 32 response output for each filter (32 x 32 x 170). Layer 3 consisted of 170 3 x 3 filters (3 x 3 x 170). Each filter was convolved with the output from layer 2 with a stride of one pixel and a pooling stride of one pixel, resulting in a 32 x 32 response output for each filter (32 x 32 x 170). Finally, layer 4 consisted of 150 3 x 3 filters (3 x 3 x 150). Each filter was convolved with the output from layer 3 with a stride of one pixel and a pooling stride of one pixel, resulting in a 32 x 32 response output for each filter (32 x 32 x 170).

The number of filters for each layer was set through preliminary testing to ensure that the overall network accuracy for discriminating between the two tasks was reasonably high (at or above 90%).

### Fully-Connected Layer and Decision Node

The fully-connected layer consisted of 280 nodes, each performing the standard weighted sum, with the output passed through a standard sigmoid activation function. The number of fully connected nodes was determined by preliminary testing. The decision layer consisted of a single node, using the same weighted sum and activation function as nodes in the fully-connected layer. Because the network made a binary decision (object task or function task), only a single output node was necessary.

### Backpropagation and Deconvolution

Backpropagation for the network followed the typical CNN operations with a few exceptions based on the simplicity of the network, its task, and to avoid over-correcting filter weights when the fit was good between any given filter and the neural response. For example, the decision layer error, *E*, simply consisted of the difference between the target response (1 = function task, 0 = object task) and the output of the decision node sigmoid. Each layer had a different learning rate, ε, to help differentially control the weight adjustments. The specific learning rates that we used were determined through preliminary testing. To update the weights in the fully-connected layer, we used the first derivative of the decision node sigmoid, the output from layer 4, and the first derivative of each fully-connected node’s sigmoid to adjust the weights in the fully-connected layer. This process computed the loss, *L,* for the layer. Updating the fully-connected layer weights then followed the usual form: *w_i_ = w_i_ + ε(L*E)*. Updating the weights in the convolutional layers followed the traditional operations (e.g., [44]), with the gradient loss from the previous layer being convolved by 180°-rotated filters from the subsequent layer, and then scaled by the network error along with the first derivative of the output node’s sigmoid to compute the gradient loss for the current layer. The weight adjustments consisted of scaling the current layer’s weights by the gradient loss along with that layer’s input. To prevent any one filter’s weights from being over-corrected due to a good fit with the neural response despite a classification error, the weight update values were further scaled by the average absolute distance between that filter’s response and the neural response. This meant that if a filter’s response was, on average, close to the neural response, the updates were reduced in proportion to these filter response distances, avoiding excessive corrections. An additional scalar was added based on the distance between each filter’s response (location specific) and the neural response. Adding that scalar only improved network performance by a fraction of a percent overall and could be removed. We chose to use it only to maximize the best possible performance, even if the difference was negligible. Finally, to help control for gradient explosion, the updated weights were L2-normalized.

To reconstruct the activation maps for each layer, image, and task, we followed the deconvolution routine of [43]. Briefly, on the forward pass of a given image and corresponding neural responses, each *trained* convolutional layer of the brain-guided network was linked to a “switch” operator that un-pooled the pooled matrix, then passed through the leaky ReLu activation function and filtered with the transpose of that layer’s filters. Then, within each layer, each filter’s activation reconstruction was summed with the other reconstructions and normalized by the maximum activation for each image.

### Network Training

The brain-guided CNN was trained stochastically, whereby each image was paired with one of two neural responses (the corresponding participant-averaged responses for the object or function task). Thus, for each training iteration, 128 images (64 paired with object task neural data and 64 paired with the function task neural data) were passed through the network. For each iteration, image order was shuffled to ensure that the network did not learn any particular pattern of input images. Training was halted when the network reached ∼90% task classification accuracy (at which point the error curve largely leveled off). We opted to use participant-averaged ERPs due to the impracticality of repeated training runs for each participant (training this particular network took a considerable amount of time for each training run). However, we found that each participant’s data were excellent predictors of the averaged set, suggesting that the mean data sets were well representative of each participant’s own data set (see **Supplementary Figure S2**). The code for the network and training routine is available upon request from the corresponding author.

### Building the Behavioral Information Utility (IU) Maps

Fifty observers were recruited on Prolific. Each observer viewed each of the 80 images and were tasked to click with a mouse or track pad the 10 image locations that were most relevant to the function or object. Each participant completed each task with images and tasks presented in a random order. Each image had an assigned function and object (see the *Experimental Procedure* section) that was written above the scene image. Observers were limited to desktop and laptop environments for the experiment (i.e., tablet, phone, and other mobile OS were blocked) to ensure that images were displayed large enough to perform the task well. Images were programmed to be displayed at 50% the height of each observer’s screen, but because each observer completed the experiment on their own computer, the actual image size varied across observers. The experiment took an average of 42 minutes to complete and observers were compensated $12 for their time.

Using the crowd-sourced click data, IU maps were built for each stimulus and task (two sets of IU maps were constructed). For each image and task, the clicks were aggregated across participants onto one map. The aggregated clicks were then convolved with a 32x32 pixel Gaussian kernel and normalized to the maximum so that the values in each IU map ranged from 0 to 1. We chose a 32x32 pixel kernel so as to minimize the overlap between object and function task-relevant regions, but that was not always possible as detailed in the Results section.

## Acknowledgements

Thank you to Isabel Gephart, Tori Gobo, and Harrison Howe for their help with data collection, and to our funding sources: James S. McDonnell Foundation grant (220020430) to BCH; National Science Foundation grant (1736394) to BCH and MRG.

## Data Availability Statement

All EEG data files, IU maps, and network code are available upon request from the corresponding author.

