## Supplementary Material for "Brain-Guided Convolutional Neural Networks Reveal Task-Specific Representations in Scene Processing"

**Supplemental Material**

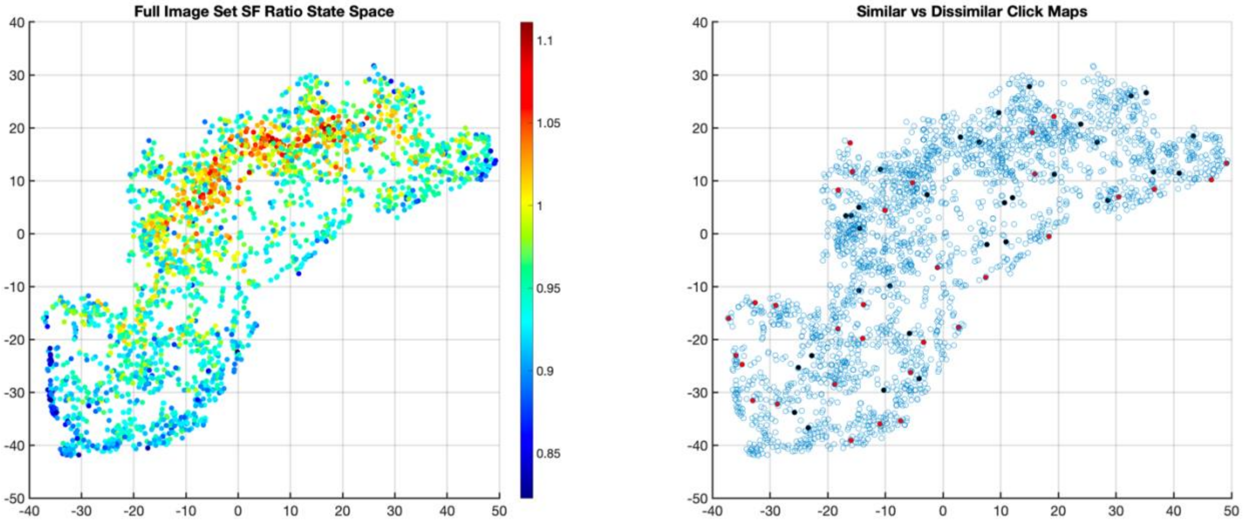


**Figure S1.** To examine whether the differences that we observed in **Figure 7** could be explained by image structure, we plotted all images from our 2500 image database in tSNE coordinates (see Method section for further detail), shown in **Figure S1**. **Figure S1a** plots each of our images as a color-coded point, with the color indicating the extent to which the image was biased toward higher or lower spatial frequencies. Specifically, each image was filtered with log-Gabor filters, each tuned to one of 9 spatial frequencies. We then summed the filter response power across the four lowest spatial frequencies and the four highest spatial frequencies and took the ratio of the two (values above 1 indicate a high spatial frequency bias). The result shows that the tSNE image space is high frequency biased in the central region, becoming increasingly low spatial frequency biased moving away in all directions from that central region. **Figure S1b** shows the same space but with the 64 images that were used for analysis in this study highlighted. Points highlighted in black are images that yielded similar IU maps, with the red points highlighting images that yielded dissimilar IU maps. No apparent clustering of images with similar vs dissimilar IU maps is observed. It therefore does not seem that the differences in ERP similarity between the two tasks can be accounted for by low-level image characteristics.

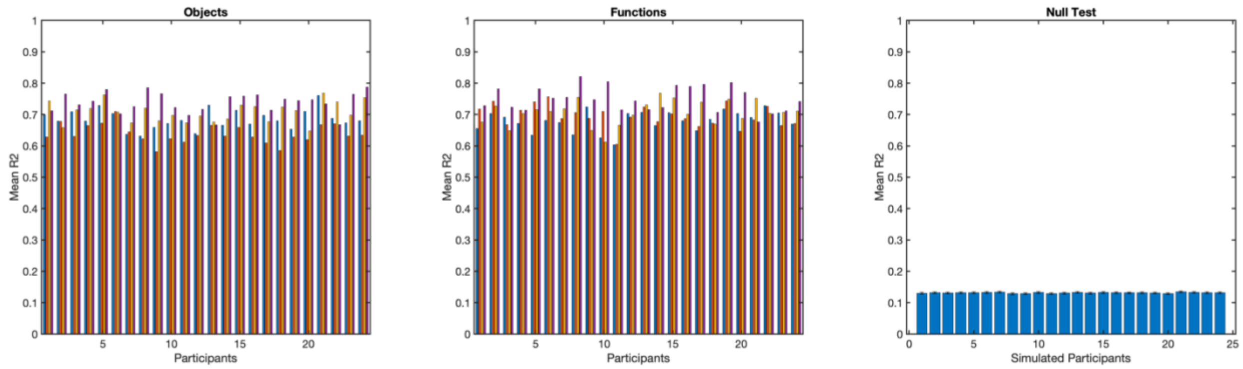


**Figure S2.** Results from the similarity analysis between each participants ERP data set (object task left, function task middle) and the group average for each task. Each set of four bars comes from each of the 24 participants. Each bar shows the electrode averaged R2 for each of the target time points within the same time 11 ms window that the network used to differentiate tasks. To the far right is what would be expected by chance by running the same similarity analysis on random data sets. Each participant's ERP data are thus well explained by the group average.
